# Cortical tracking of speech is reduced in adults who stutter when listening for speaking

**DOI:** 10.1101/2024.02.23.581767

**Authors:** Simone Gastaldon, Pierpaolo Busan, Nicola Molinaro, Mikel Lizarazu

**Author notes:** **Corresponding author:** Simone Gastaldon Dipartimento di Psicologia dello Sviluppo e della Socializzazione (DPSS) Università di Padova Via Venezia 8, 35131, Padova (PD), Italy.

## Abstract

**Purpose:** Investigate cortical tracking of speech (CTS) in adults who stutter (AWS) compared to typically fluent adults (TFA) to test the involvement of the speech-motor network in tracking auditory information.

**Method:** Participants’ EEG was recorded while they either had to simply listen to sentences (listening only) or complete them by naming a picture (listening-for-speaking), thus manipulating the upcoming involvement of speech production. We analyzed speech-brain coherence and brain connectivity during listening.

**Results:** During the listening-for-speaking task, AWS exhibited reduced CTS in the 3-5 Hz range (theta), corresponding to the syllabic rhythm. The effect was localized in left inferior parietal and right pre-/supplementary motor regions. Connectivity analyses revealed that TFA had stronger information transfer in the theta range in both tasks in fronto-temporo-parietal regions. When considering the whole sample of participants, increased connectivity from the right superior temporal cortex to the left sensorimotor cortex was correlated with faster naming times in the listening-for-speaking task.

**Conclusions:** Atypical speech-motor functioning in stuttering also impacts speech perception, especially in situations requiring articulatory alertness. The involvement of frontal and (pre-) motor regions in CTS in typically fluent adults is highlighted. Speech perception in individuals with speech-motor deficits should be further investigated, especially when smooth transitioning between listening and speaking is required, such as in real-life conversational settings.

## Introduction

Developmental Stuttering (DS, also known as Childhood-Onset Fluency Disorder; American Psychiatric Association, 2013) is a neurodevelopmental disorder of the normal flow of speech, characterized by symptoms such as blocks, prolongations, and repetitions. People who stutter know what they want to say, but they may be unable to speak in a fluent manner. Importantly, DS may persist in adulthood, impairing the quality of life of affected individuals (Craig et al., 2009; Nang et al., 2018).

DS likely has a multifactorial origin, comprising motor, linguistic, emotional, neural and genetic factors (Smith & Weber, 2017). In particular, in recent years, genetic factors have been identified (Barnes et al., 2016; Benito-Aragón et al., 2020; Chow et al., 2020; Frigerio-Domingues & Drayna, 2017; Kang et al., 2010; Kang & Drayna, 2012; Kraft & Yairi, 2011), which may facilitate the appearance of a series of atypical characteristics of the central nervous system (Alm, 2021a; Craig-McQuaide et al., 2014; A. C. Etchell et al., 2018). In this context, affected neural functions may result in impaired capacities related to the sensorimotor planning and execution of speech (Alm, 2021b; Chang et al., 2019), which in turn is reflected in altered sensorimotor brain rhythms (Etchell et al., 2016; Ghaderi et al., 2018; Jenson et al., 2018, 2020; Joos et al., 2014; Saltuklaroglu et al., 2017). DS seems to be characterized by the presence of a deficit in internal timing and motor coordination (Alm, 2004), involving wide neural systems and comprising regions such as the basal ganglia, the supplementary motor area, the inferior frontal cortex, and temporal regions (Busan, 2020; Busan et al., 2019; Craig-McQuaide et al., 2014; Etchell et al., 2018; Watkins et al., 2008). Importantly, neural inefficiency that perturbs speech-motor execution seems to also affect aspects of speech comprehension, specifically leading to weaker or less efficient predictive processing (Gastaldon et al., 2023). Similarly, audio-motor interactions appear to be bi-directionally impaired in DS, with deficits in properly integrating auditory feedback in speech production (Bradshaw et al., 2021; Chang et al., 2016; Daliri & Max, 2015, 2018; Halag-Milo et al., 2016; Hesse, 2023; Kim et al., 2020). From a behavioral point of view, conflicting results have been reported, with some studies finding malfunctioning of predictive timing during auditory-motor coupling in people who stutter can lead to some differences when compared to fluent speakers (Falk et al., 2015) and others reporting no overt differences in auditory-motor integration (Assaneo et al., 2022). Therefore, audio-motor interaction seems instrumental for effective speech production and to be inefficient in DS.

However, evidence suggests that such interaction is also relevant for aspects of speech perception, specifically in the temporal alignment of internal low frequency brain rhythms (delta – 0.1-3 Hz and theta – 4-7 Hz – frequency bands) to acoustic energy fluctuations (envelope) of the external speech signal. This phenomenon is known as cortical tracking of speech (CTS), sometimes also referred to as “speech-brain entrainment”, and is often considered to be a valuable index reflecting the efficiency of neural processing of quasi-rhythmic components of speech, specifically of prosodic (delta) and syllabic (theta) information (Assaneo & Poeppel, 2018; Molinaro & Lizarazu, 2018; Poeppel & Assaneo, 2020; Poeppel & Teng, 2020). Importantly, a growing body of evidence supports the view that frontal, motor and premotor regions modulate CTS in the auditory cortex in a top-down manner (Keitel et al., 2018; Park et al., 2015). Evidence also suggests that there is a preferred frequency range at which motor and auditory cortices are coupled, i.e., within the theta range between 3 and 5 Hz, with a peak at 4.5 Hz, (Assaneo & Poeppel, 2018), a range associated with production and perception of syllables across languages (Ding et al., 2017, Poeppel & Assaneo, 2020). In this scenario, the motor system is supposedly exploited to reduce noise and uncertainty, by generating temporal predictions via efferent motor signals, causing phase-resetting in auditory cortices and optimizing sensory perception (Rimmele et al., 2018). Recent behavioral evidence supports this account: higher individual speech production rates and auditory-motor coupling were associated with better performance in a speech comprehension task (Lubinus et al., 2023).

Importantly, altered brain processes related to CTS have been proposed as a risk factor for the appearance of developmental speech and/or language disorders, usually characterized by abnormalities in cortico-basal-thalamo-cortical circuitries involved in the processing of sensory cues (such as beats in music and/or linguistic meter in speech), thus playing a role in processing and predicting events in a sequence (Ladányi et al., 2020). For example, CTS is differently modulated in people with dyslexia (Lizarazu et al., 2015; Molinaro et al., 2016; Lizarazu et al., 2021a). This may also be the case for people who stutter, especially if auditory-motor coupling is a contributory factor to both DS and CTS. Crucially, no evidence is currently available for brain processes related to CTS in stuttering. Thus, a better understanding of these phenomena in DS should be useful for improving our comprehension of 1) neuro-pathological mechanisms related to stuttering (both in the contexts of speech production and perception), and 2) the neural mechanisms involved in typical speech perception and production (and in their possible mutual interactions).

To address both these issues, in the present study we investigated whether adults who stutter (AWS; stuttering onset during childhood and persisting into adulthood) show altered tracking of the speech signal when compared to typically fluent adults (TFA; no diagnosis of speech disorders). Specifically, we measured speech-brain coherence on electroencephalographic (EEG) data, both at the sensor and the neural source level, during sentence listening in conditions that either overtly recruited the articulatory system (completing the sentence by naming a picture; listening-for-speaking) or not (listening to the entire sentence; listening only), in order to assess whether the upcoming involvement of the speech-motor network may have modulatory effects on CTS (see Figure 1 and Materials and Methods). While not directly simulating everyday dyadic conversations, the listening-for-speaking task still implies an alertness of the speech-motor system, in addition to higher level processes such as anticipation and planning (Corps et al., 2018), similar to the demands of conversational and turn-taking settings. By following Assaneo & Poeppel (2018), we expected to find group differences in a restricted range within the theta band, at which activity in the auditory and motor regions is supposed to be inherently coupled, reflecting tracking of syllabic rhythm.

**Figure 1.**
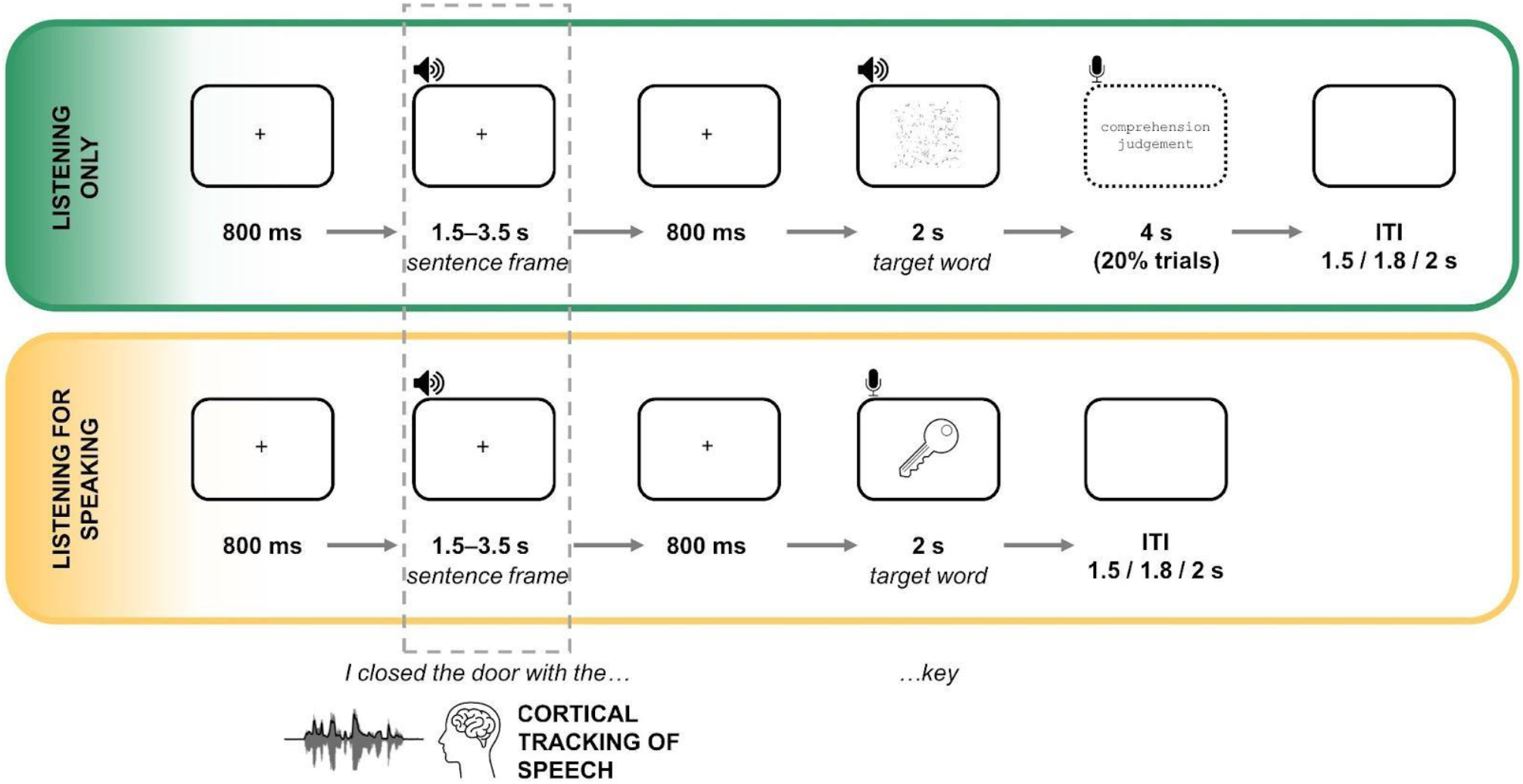
Experimental design. Participants listened to sentence frames and target final words in the listening-only task (with occasional true/false judgment questions), while they had to complete the sentence frame by naming a picture (representing the target word) in the listening-for-speaking task. Indexes related to cortical tracking of speech and cortical connectivity were measured during the auditory presentation of the sentence frames. Response times in the listening-for-speaking task were collected (calculated at picture onset).

In addition, partial directed coherence (PDC) was analyzed to quantify directional neural interactions between brain regions implicated in sensorimotor speech processing. This analysis allows us to investigate frequency-specific directional communication between cortical regions during envelope tracking, possibly highlighting differences in speech-relevant pathways. Based on Hickok and Poeppel’s dual route model of speech processing (Hickok & Poeppel, 2004, 2007), we hypothesize that neural connectivity is reduced in stuttering especially in the dorsal stream, suggested to be responsible for motor-auditory transformations (Hickok et al., 2011). This hypothesis is also supported by evidence showing that white matter tracts considered to be part of the dorsal stream are usually altered in people who stutter (Kronfeld-Duenias et al., 2016; Neef et al., 2018a, 2022; Sommer et al., 2002; Watkins et al., 2008). However, we remain agnostic as to specific patterns (directionality) and additional group differences (potential stronger connectivity in AWS in other pathways, reflecting compensatory mechanisms). To this extent, the PDC analysis is partially theoretically-driven (identify pathways compatible with a dorsal processing stream) and partially exploratory.

Thus, given the picture outlined above, we can hypothesize that:

1. CTS may be reduced in AWS relative to TFA, regardless of the listening condition (listening only vs listening-for-speaking). Alternatively, differences may be detected only when listening is coupled with the upcoming necessity to overtly activate the speech-motor system, which is consequently kept in an “alert mode” in order to appropriately initiate speech (see “Sensor level analysis” section).
2. In AWS, reduced CTS may be found in speech-motor and premotor regions, in addition to auditory and associative regions (see “Source level analysis” section).
3. In AWS, regions that are considered to be part of the dorsal stream (inferior frontal cortex, premotor and supplementary motor regions, sensorimotor and temporo-parietal regions) may be communicating less efficiently with auditory regions during speech tracking, thus displaying reduced connectivity (see “Partial directed coherence (PDC) analysis”).

## Materials and Methods

### Participants

We analyzed CTS in a dataset collected for a previous study on spoken sentence processing in adults who stutter which focused on different time-windows, used different analyses and had different aims (Gastaldon et al., 2023). The original study included 14 right-handed male adults who stutter (AWS) and 14 right-handed typically fluent male adults (TFA). The participants were matched for age and handedness. All participants were native speakers of Italian. The original study was approved by the Ethical Committee for Psychological Research of the University of Padova and conducted in accordance with the Declaration of Helsinki. We refer the reader to the original study for further details on AWS recruitment and assessment. Out of the 28 participants of the original study, four participants were excluded due to excessively noisy EEG data for the present type of analysis during sentence frame presentation for the analyses conducted here. The remaining 24 participants, 12 AWS (mean age = 34.44, SD = 9.37) and 12 TFA (mean age = 33.42, SD = 8.94), were also matched for handedness, as assessed by means of the Edinburgh Handedness Inventory (Oldfield, 1971): AWS: mean = 83.75, SD = 20.57; TFA: mean = 85.00, SD = 23.06. From the original study we also retrieved data about the Stuttering Severity Index (SSI-4; Riley, 2009) of each AWS: the higher the SSI-4 score, the more severe the stuttering. The final set was composed of 6 participants with very mild severity, 4 with mild and 2 with severe. Exploratory correlations with CTS and connectivity data were performed (statistical threshold for explorative correlations: p ≤ 0.01, “two-tailed”).

### Stimuli and procedure

The stimuli were the same used in Gastaldon et al. (2020, 2023) (see the OSF repository for additional information on stimuli characteristics: https://osf.io/tcbsh/). They consisted of 256 sentence frames (sentences without the final word, ranging in duration from 1.55 to 3.54 s; mean duration = 2.39, SD = 0.4), which were paired with 128 target words and 128 b/w line pictures (124×124 pixels), such that each word and picture appeared twice, completing a high and low constraint sentence frame. Spoken stimuli were uttered by a female native Italian speaker, recorded and digitized at 44.1 kHz using Audacity®. Audio files (*.wav) were also segmented using Audacity. During the task, participants listened to the sentence frames, then, after an 800 ms pause, either they heard a word (listening-only task) or had to produce it by naming a picture (listening-for-speaking task), in two distinct blocks, which were counterbalanced across participants. In the listening-only task, true/false comprehension questions were asked at the end of the trial in 20% of the trials, to maintain the participant’s engagement (see also Figure 1). Due to the aims of the original study (Gastaldon et al., 2023), half the sentence frames induced highly constraining contexts for the final word and half induced weakly constraining contexts. However, in order to allow for a better estimation of CTS and increase signal-to-noise ratio (SNR) and statistical power, we did not divide the sentence frames into high vs low constraining contexts. We recognize that this may be a highly relevant variable that should be investigated in future studies (for a study in the normal population, see Molinaro et al., 2021); however, here we were limited in terms of SNR and number of trials. Therefore, here we focused on the manipulation of task demands, which implied two different listening conditions: listening for comprehension (listening only), or listening in order to complete the sentence as quickly as possible by naming a picture (listening-for-speaking). Participants sat in a dimly lit room in front of a computer screen. The experimental material was delivered through E-Prime 2.0 (Psychology Software Tools, Pittsburgh, PA). Auditory stimuli were presented through built-in speakers. Responses (picture naming and true/false answers) were collected via a microphone set in front of the participant. In the listening-for-speaking task, audio recording started at the onset of the picture to be named and lasted for 2 seconds. The experimental paradigm is exemplified in Figure 1. For further details on the experimental design, we refer the reader to the original study (Gastaldon et al., 2023).

### EEG data acquisition and preprocessing

During the task, the electroencephalogram (EEG) was recorded using a BrainAmp amplifier and BrainVision Recorder software (BrainProducts, Germany). EEG was recorded using 64 electrodes that were positioned according to the international 10-10 system (Nuwer et al., 1998). Scalp-electrode impedance was kept below 10Ω. The recording was referenced to the left earlobe. Electrode AFz served as the ground. Two electrodes at the outer canthi of both eyes recorded horizontal eye movements and one electrode below the left eye recorded vertical eye movements. EEG was sampled at 1000 Hz and band-pass filtered online from 0.1 to 1000 Hz.

The preprocessing pipeline for the present work was the following. Heartbeat and EOG artifacts were identified using independent component analysis (ICA) and subtracted from the recordings in a linear manner. The ICA decomposition was carried out using the Infomax algorithm implemented in the Fieldtrip toolbox (Oostenveld et al., 2011). Across participants, the number of heartbeat and ocular components that were removed varied from 1 to 4 and 1 to 3 components, respectively. Furthermore, trials were visually inspected to discard any remaining artifacts. Bad channels were substituted with interpolated values computed as the average of the neighboring electrodes obtained through the triangulation method implemented in Fieldtrip. A minimum of 75% artifact-free trials per participant was required for inclusion in subsequent analyses. As noted above, this led to the exclusion of two participants from each group, resulting in a final sample of 24 participants (12 AWS and 12 TFA). In the case of TFA, an average of 4.89% (SD = 3.1) trials and 5.23% (SD = 3.8) trials were excluded for comprehension and production tasks, respectively. Similarly, for AWS, an average of 5.4% (SD = 3.59) trials and 8.13% (SD = 3.86) trials were excluded for comprehension and production tasks, respectively. Importantly, no significant group or task differences were observed in the number of excluded trials (all *T*s < 1.6, all *p*s > 0.11, two-tailed t-test). EEG data and MATLAB scripts for the analyses described in the following sections are available on a dedicated OSF repository: https://osf.io/7gpyb/.

### Cortical tracking of speech (CTS) analysis

#### Sensor level analysis

Coherence measures the degree of phase synchronization between two signals in the frequency domain. For each participant and condition, we used coherence to quantify the cortical tracking of speech (CTS), which represents the coupling between the speech temporal envelope and cortical oscillations. We obtained the envelope of the speech signal from the Hilbert transformed broadband stimulus waveform. According to previous research in speech processing we expected to find strong CTS in the low-frequency (< 10 Hz) spectrum and in temporal sensors (Molinaro et al., 2016; Molinaro & Lizarazu, 2018; Lizarazu et al., 2021b; Ershaid et al., 2024; Issa et al., 2024). Therefore, we selected a set of 12 channels, evenly distributed to cover the temporal lobes of the brain – precisely, 6 channels allocated over the left hemisphere (C3, C5, CP3, CP5, FC3, FC5) and additional 6 over the right hemisphere (C4, C6, CP4, CP6, FC4, FC6). Artifact-free trials were segmented into 1-second windows with 50% overlap. Coherence was then calculated using the cross-spectral density of the FFT of the two signals (i.e., speech envelope and EEG data segments), normalized by the power spectrum of each signal. For each EEG sensor, coherence was calculated in the 1 – 15 Hz frequency band with 1 Hz (inverse of the segment duration) frequency resolution (Molinaro et al., 2016; Molinaro & Lizarazu, 2018). This procedure was followed for each participant and task/listening condition.

To estimate the coherence bias, the auditory envelopes were randomly shuffled across epochs for each participant, and coherence was recalculated in 100 permutations. The coherence data from the selected sensors of interest were separately averaged for each hemisphere and then transformed into z-scores using the mean and standard deviation derived from the 100 random EEG-audio pairings for those sensors. For each condition and frequency bin, z-score transformations were computed using the task-specific mean and standard deviation obtained from the random pairing dataset, and with an equal number of trials as the actual EEG-audio pairing dataset.

For the statistical analysis, we calculated the mean CTS values (z-scored coherence) within the theta band, specifically in the 3-5 Hz frequency range. We focused on this frequency range because of two specific reasons: 1) a data-driven reason is that a peak is present in our auditory stimuli in the same frequency range, indicating syllabic rhythm (see Supplementary Figure 2), and 2) a theoretically-driven reason is the frequency-restricted preference for the coupling between auditory and motor regions, as explained in the Introduction (see also Assaneo & Poeppel, 2018). To assess group differences in each task, we conducted an ANOVA on the z-transformed coherence values, with hemisphere (left vs right) as the within-subject factor and group (TFA vs AWS) as the between-subject factor (considering effects of main factors and their interaction; post-hoc analyses conducted using t-test; statistical threshold at p ≤ 0.05, two-tailed).

#### Source level analysis

Coherence values were also estimated at the source level for each participant and condition in the theta band (3-5 Hz), where significant results were observed at the scalp level. For the source level analysis, we utilized a frequency-domain adaptive spatial filtering imaging of coherent sources algorithm (Gross et al., 2001), implemented in the Fieldtrip toolbox. To establish the spatial relationship between electrode positions of the participants (defined with a template electrode layout) and the cortical mesh, we employed a standard boundary element head model (BEM) extracted from the Montreal Neurological Institute (MNI) template. This BEM consists of three 3-D surfaces (skin, skull, brain) derived from the MNI dataset. The forward model was computed using an 8 mm grid encompassing the entire brain compartments of the BEM, representing various source positions. To perform source analysis, we constructed common space filters utilizing the leadfield of each grid point and the cross-spectral density matrix (CSD). The CSD matrices were computed within the theta (4 Hz with ±1 Hz frequency smoothing) band by applying the fast Fourier transform to 1-second data segments in sliding windows shifting in 0.5 seconds steps. As anticipated, the selection of the theta range was based on the observation of group effects at the sensor level occurring specifically at this frequency. Beamformer coefficients were computed considering the dominant source direction within all voxels and a regularization factor of 7% was applied. The coherence for each source location was estimated using the EEG data and the spatial filter in the theta band. To ensure comparability of source coherence values across subjects, we normalized individual coherence brain maps. For this reason, the coherence at each source was converted to a z-score value by subtracting the mean coherence across all sources and dividing by the standard deviation across all sources. Successively, for each group and condition, z-scored source coherence values were projected on the brain surface mesh image BrainMesh_ICBM152_smoothed from Surf Ice (Version 12.1; https://www.nitrc.org/projects/surfice/)

Finally, based on previous functional neural evidence on DS we selected five regions of interest (ROIs) from the Automatic Anatomical Labeling (AAL; Tzourio-Mazoyer et al., 2002). More specifically, ROIs were defined considering that stuttering mainly affects neural networks that are fundamental for sensorimotor processing, thus impairing speech planning, programming, and execution (compare with Chang et al., 2019). In this context, abnormal neural activity in areas such as the inferior frontal cortex, primary somato-motor regions, auditory cortex, supplementary motor area, premotor cortex, and associative regions (such as the parietal cortex) have been consistently reported as “neural markers” of DS (see Belyk et al., 2015, 2017; Brown et al., 2005; Budde et al., 2014; Busan, 2020; Busan et al., 2019; Chang & Guenther, 2020; Chang et al., 2019; Craig-McQuaide et al., 2014; Etchell et al., 2018; Ingham et al., 2012; Neef et al., 2015; Zhang et al., 2022). As a consequence, within each cerebral hemisphere, we defined the subsequent “clusters’’ of brain regions of interest (as shown in Figure 2): i) the inferior frontal gyrus (IFG, comprising the pars opercularis, pars triangularis, and pars orbitalis), ii) the premotor and supplementary motor cortex (preM), iii) the sensorimotor strip (SM, comprising the pre-central and post-central gyri), iv) the inferior parietal lobule (IPL, comprising the supramarginal and angular gyri), and v) the superior temporal gyrus (STG).

**Figure 2.**
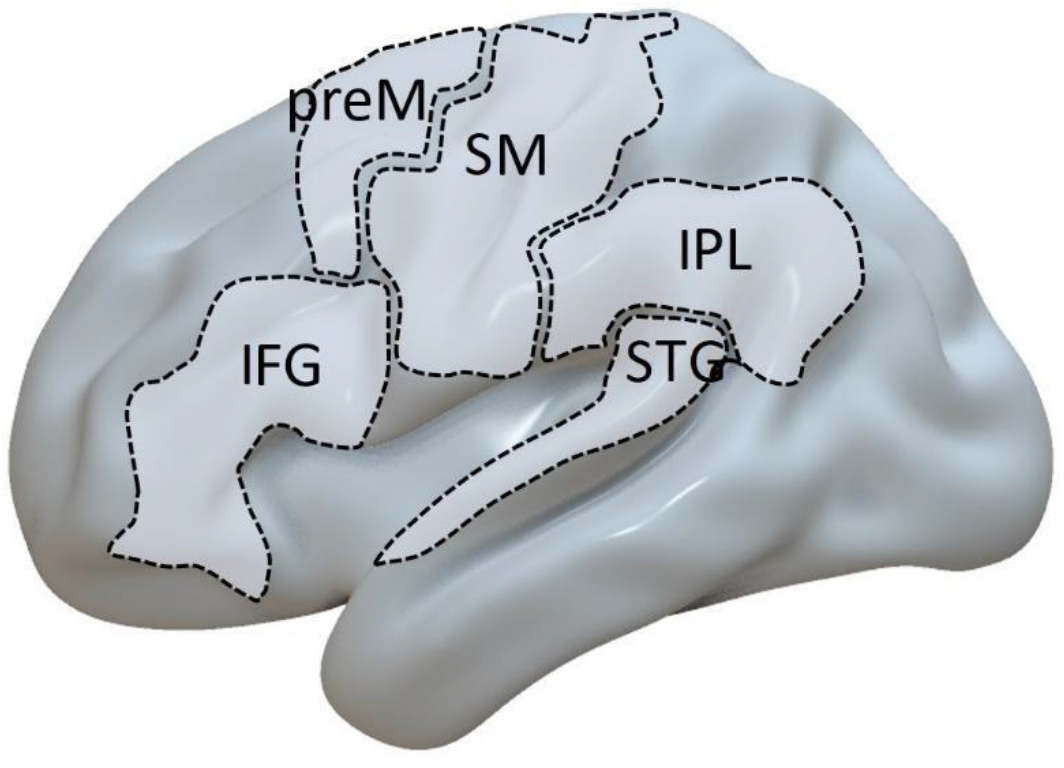
Regions of interest (ROIs) selected for statistical comparison between groups in the source and connectivity analyses. Five ROIs were selected in the left and right hemisphere: i) the inferior frontal gyrus (IFG, comprising the pars opercularis, triangularis, and orbitalis), ii) the premotor and supplementary motor cortex (preM), iii) the somato-motor strip (SM), iv) the inferior parietal lobule (IPL), and v) the superior temporal gyrus (STG).

For each task, we employed the Wilcoxon ranked sum non-parametric test to assess group differences on the mean of the z-scored coherence values within each ROI (statistical threshold at p ≤ 0.05, two-tailed).

#### Partial directed coherence (PDC) analysis

We employed partial directed coherence (PDC) to assess the causal connections between neural activity associated with speech processing within our designated ROIs (IFG, preM, SM, IPL and STG). After creating spatial filters, virtual time series in the source locations within the ROIs were reconstructed by applying the respective spatial filter to the EEG sensor data filtered in the theta (3 - 5 Hz) band. Because ROIs typically comprise many point sources, we employed principal component analysis (PCA) to identify the most representative time series within each ROI. To achieve this, we conducted a PCA on all time-series within each ROI and selected the first principal vector, which represented the distribution that explained most of the variance across all time-series that entered the PCA. For each participant and task, we computed PDC between the representative time series in each ROI. PDC is based on the Granger Causality principle (Granger, 1969; Seth et al., 2015) and on vector autoregressive (VAR) modeling of the data. The VAR model of order *p* for a variable *x* is given by:

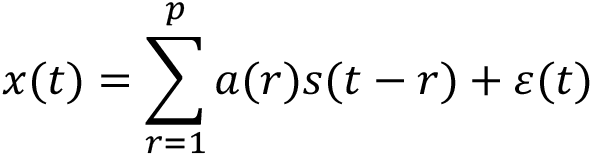

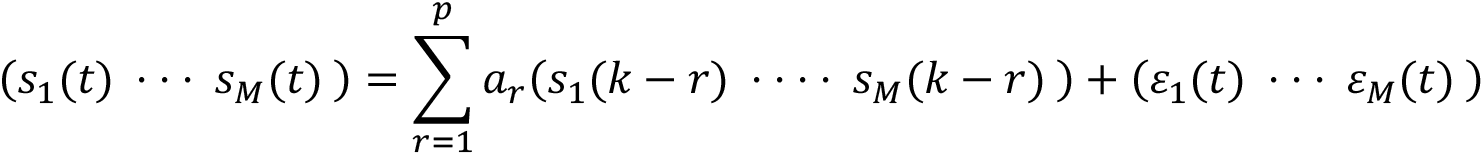

where s(t) = (s_1_(t), s_2_(t), …, s_M_(t)) are the stationary M-dimensional simultaneously measured time series in each ROI; ar are the M x M coefficient matrices of the model; and (t) is a multivariate Gaussian white noise process. In our case, M = 10 since we calculated the connectivity network formed by five different ROIs. The model order *p* was selected with the Schwartz Information Criterion. This criterion selects the model order that optimizes the goodness of fit of the model, while introducing a penalty depending on the complexity of the model. In the frequency domain the version of Granger-causality is given by:

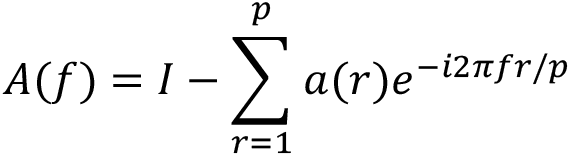

The first term of the difference refers to the identity matrix (M-dimensional) and the second one to the Fourier transform of the VAR coefficients. Then, the PDC from the ROI j to ROI i is given by:

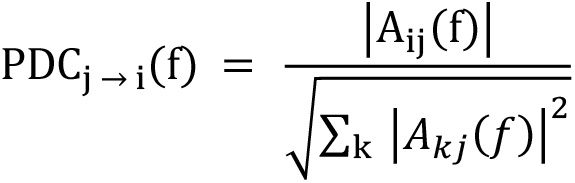

The PDC provides a measure of the linear directional coupling strength of *s_j_* on *s_i_* at frequency *f* (theta). The PDC values vary between 0 (no directional coupling) and 1 (perfect directional coupling). PDC analysis was performed using the Frequency-Domain Multivariate Analysis toolbox (FDMa, Freiburg Center for Data Analysis and University of Freiburg, Germany), and the model order was computed using algorithms developed in the Multivariate Autoregressive Model Fitting (ARfit) software package (Schneider & Neumaier, 2001). To assess group differences, separately for each task we used the Wilcoxon ranked sum non-parametric test on PDC values (statistical threshold at p ≤ 0.05, two-tailed).

### Naming accuracy and response times analysis (listening-for-speaking task)

For naming latencies (response times, RTs), we took the data from Gastaldon et al. (2023), also available here: https://osf.io/5jkur/. Here we summarize how latencies were derived in the original study, but we refer the reader to the original article for additional details. To estimate naming times, audio recordings (2 seconds *.wav files starting at picture onset) were fed to Chronset (Roux et al., 2017). Only correct responses were considered. Responses were coded as incorrect if they started with hesitation sounds, if corrections were made during the response, or if the participant could not produce enough of the target word in the 2-second recording (in order to be able to assess the correctness of the response).

Statistical analyses were performed in R. Accuracy was analyzed with a generalized linear mixed-effects model (GLMM) with binomial distribution family. Group, lexical frequency of the target word (retrieved through PhonItalia; Goslin et al., 2014) and repetition (the same target picture was presented twice in the task, associated with two different sentence frames) were set as fixed effects, while participant and item as random intercepts. RTs were analyzed with a GLMM with gamma distribution family and identity link function. Group, lexical frequency of the target word and repetition were set as fixed effects, while participant and item as random intercepts. As explained above, we decided not to include sentence constraint as a factor here since for the coherence analysis (the main focus of the present work) we did not differentiate between the two conditions for methodological reasons. GLMM were fitted with the *lme4* package (Bates et al., 2015) and contrasts set to sum coding. Finally, as for SSI-4, RTs were correlated with CTS and connectivity data (statistical threshold at p ≤ 0.01, two-tailed).

## Results

### Naming (listening-for-speaking task)

Accuracy and response times (RTs) are shown in Figure 3, while model summaries are reported in Table 1. Participants of both groups had a very high accuracy in producing the correct word (AWS: mean = 0.96, SD = 0.2; TFA: mean = 0.99, SD = 0.1; see Figure 3A). However, the model revealed a main effect of repetition (higher accuracy when the picture appeared for the second time) and a main effect of group, with AWS less accurate than TFA (see Table 1). Regarding response times, AWS were slower than TFA (AWS: mean = 771.19 ms, SD = 267.77; TFA: mean = 650.53 ms, SD = 219.98; see Figure 3B). The model revealed a main effect of repetition and, importantly, a main effect of group (see Table 1). To test the robustness of the results to possible outliers for accuracy, we re-run the analysis by excluding the AWS participant with accuracy = 0.84 (see Figure 3), and the results are still consistent (main effect of group: t = -2.35, p = 0.019, 95%CI [0.33 - 0.91]). Speculatively, lower accuracy, rather than reflecting possible inefficiency in retrieving lexical items in AWS, is likely due to the limited time available for recording the responses (2 seconds after picture onset): sometimes, AWS may have provided the correct response outside this window, making it impossible to evaluate their response off-line, hence the reduced accuracy (i.e., fewer trials coded as correct). This interpretation is compatible with the fact that the accuracy outlier in the AWS group (accuracy = 0.84) is also the one with longest mean RTs for correct responses (RT = 1155 ms).

**Figure 3.**
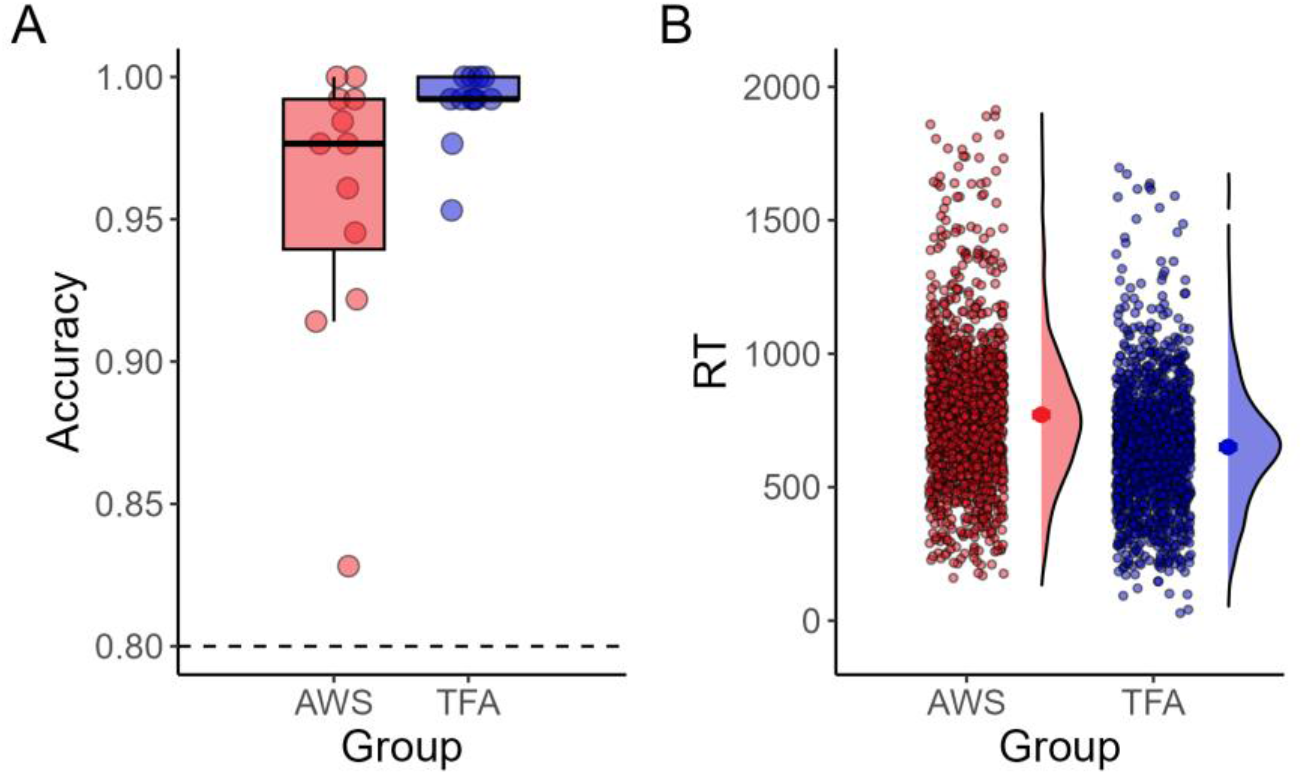
Behavioral results for the listening-for-speaking task. A) Subject-level (individual) accuracy scores (dots) and boxplots; note: y-axis starting at 0.8. B) Single-trial response times (dots), group-level means with error bars and density distributions.

**Table 1.** Model summaries for accuracy and RTs for naming in the listening-for-speaking task.

| ACCURACY |  |  |  |  |
| --- | --- | --- | --- | --- |
| Predictor | Estimate | 95% CI | Statistics | p-value |
| (Intercept) | 60.83 | 20.21 – 183.11 | 7.31 | <0.001 |
| Lexical frequency | 1.22 | 0.97 – 1.54 | 1.68 | 0.093 |
| Group | 0.47 | 0.27 – 0.82 | -2.64 | 0.008 |
| Repetition | 0.70 | 0.55 – 0.90 | -2.84 | 0.005 |
| RESPONSE TIMES |  |  |  |  |
| Predictor | Estimate | 95% CI | Statistics | p-value |
| (Intercept) | 730.15 | 688.82 – 771.48 | 36.64 | <0.001 |
| Lexical frequency | -6.05 | -14.21 – 2.10 | -1.45 | 0.146 |
| Group | 60.43 | 34.44 – 86.41 | 4.56 | <0.001 |
| Repetition | 36.44 | 29.12 – 43.76 | 9.76 | <0.001 |

### Sensor-level CTS

Initially, we conducted an assessment of sensor-level cortical tracking of speech within the 1 - 15 Hz frequency range for each group (TFA and AWS) and task (listening-only and listening-for-speaking). Consistent with previous studies, we observed that during speech listening, CTS was highest in the theta (3-5 Hz) frequency band (Figure 4A) in bilateral fronto-central, central, and centro-parietal sensors (Figure 4B), consistent with the topography usually found in M/EEG studies on coherence as measure of CTS in the theta range (Destoky et al., 2019).

**Figure 4.**
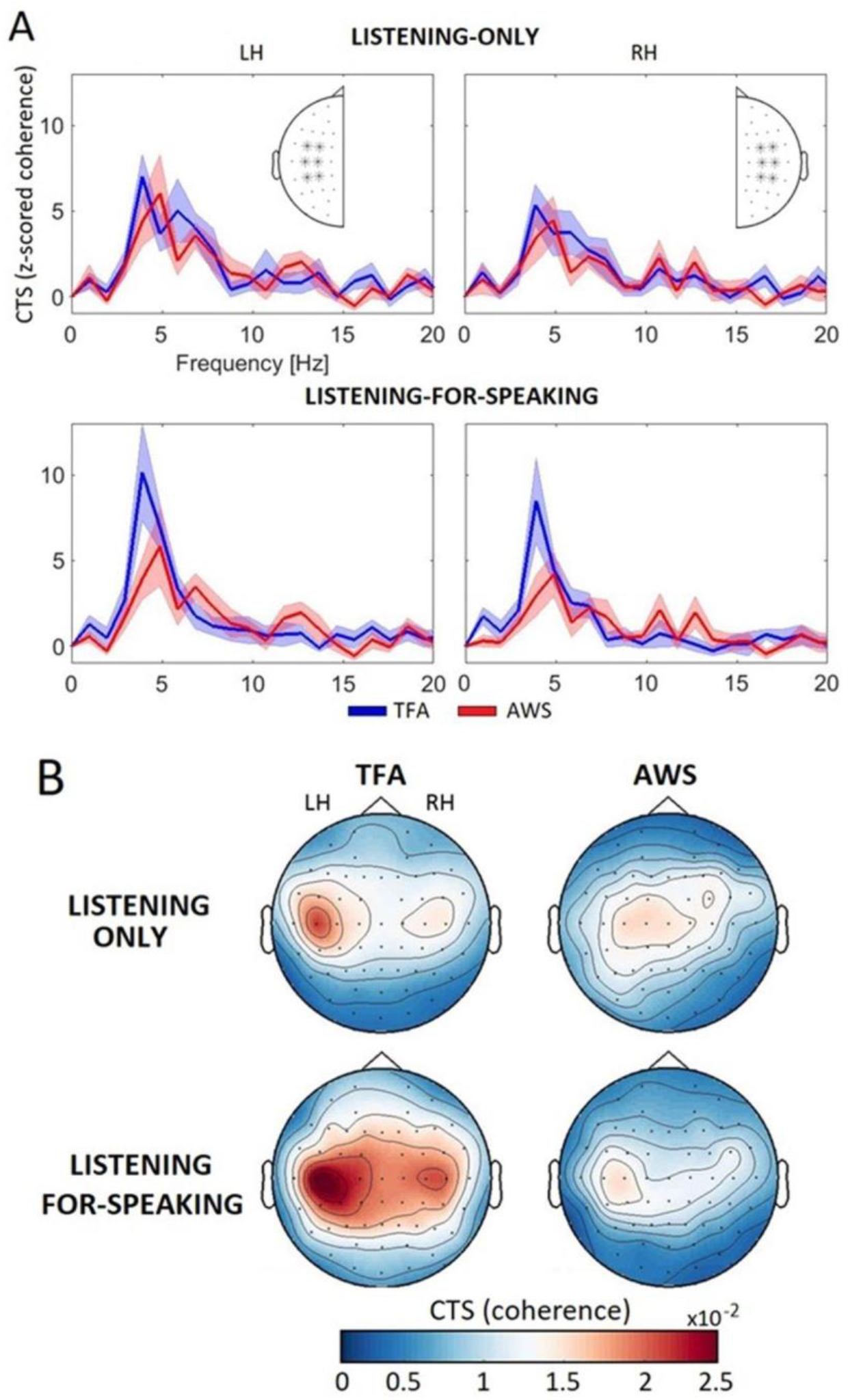
Spectra distribution and topographic map of the CTS at the sensor level. A) Corrected coherence values (coherence values converted into z-scores using the mean and standard deviation derived from the 100 random EEG-audio combinations) in the 1 – 15 Hz frequency range can be observed across representative sensors (C3, C5, CP3, CP5, FC3, FC5, C4, C6, CP4, CP6, FC4, FC6) of the left (LH) and right (RH) hemisphere. B) For each group (TFA: Typical Fluent Adults; AWS: Adults Who Stutter) and task (listening-only and listening-for-speaking), we plotted the topographic maps of uncorrected coherence values in the theta (3 - 5 Hz) frequency band.

For each task, we performed an ANOVA on the mean CTS values (z-scored coherence) within the theta band and across the sensors of interest in both the left and right hemispheres. In the listening-only task, we did not observe any main effects or interactions in the CTS values (all *F*(1,22) < 1.97, all *p* < 0.17, *ղ2* <0.06). However, we did observe a main effect of Group (*F*(1,22) = 4.07, *p* = 0.05, *ղ2* = 0.15) in the CTS values for the listening-for-speaking task. Post-hoc tests showed that CTS was significantly higher in TFA compared to AWS (*t* = 2.02, *p* = 0.05, Cohen’s d = 0.80). No statistically significant correlations with RTs or SSI-4 were found.

### Source-level CTS

When considering source analyses, we observed that for both the listening-only and the listening-for-speaking tasks, frontal, temporal, and parietal cortical regions showed strong CTS (z-scored coherence values) in the theta band (Figure 5). Subsequently, we calculated the mean of the CTS values in each of the ROIs described in the Materials and Methods section: the inferior frontal gyrus (IFG), the premotor/supplementary motor cortex (preM), the sensorimotor strip (SM), the inferior parietal lobule (IPL), and the superior temporal gyrus (STG).

**Figure 5.**
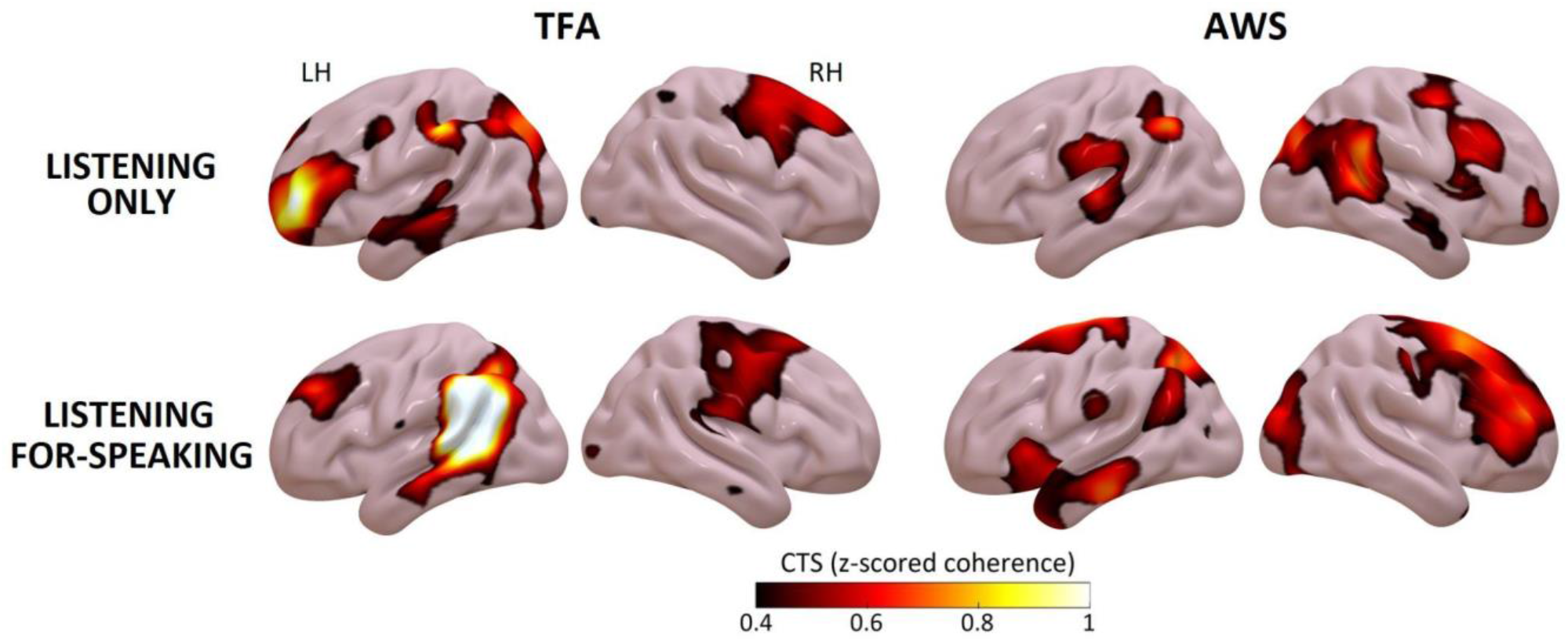
Source reconstruction of the CTS values in the theta range. For each group (TFA: Typical Fluent Adults; AWS: Adults Who Stutter) and listening condition (listening only and listening-for-speaking), we plotted the source maps of CTS values (coherence values converted into z-scores using the mean and standard deviation derived from the CTS values in all the sources) in the theta (3 - 5 Hz) frequency band.

In line with the results observed at the sensor level, we found that the CTS values were significantly stronger for individuals with TFA compared to AWS in the left IPL (MTFA = 1.92, SDTFA = 2.58; MAWS = 0.22, SDAWS = 0.34; *p* = 0.03) and in the right preM regions (MTFA = 0.07, SDTFA = 1.21; MAWS = -0.74, SDAWS = 0.36; *p* = 0.01), only for the listening-for-speaking task (Figure 6). We did not find any group differences in the listening-only task (all *p* > 0.09) (Supplementary Figure 1). No statistically significant correlations with RTs and SSI-4 were found.

**Figure 6.**
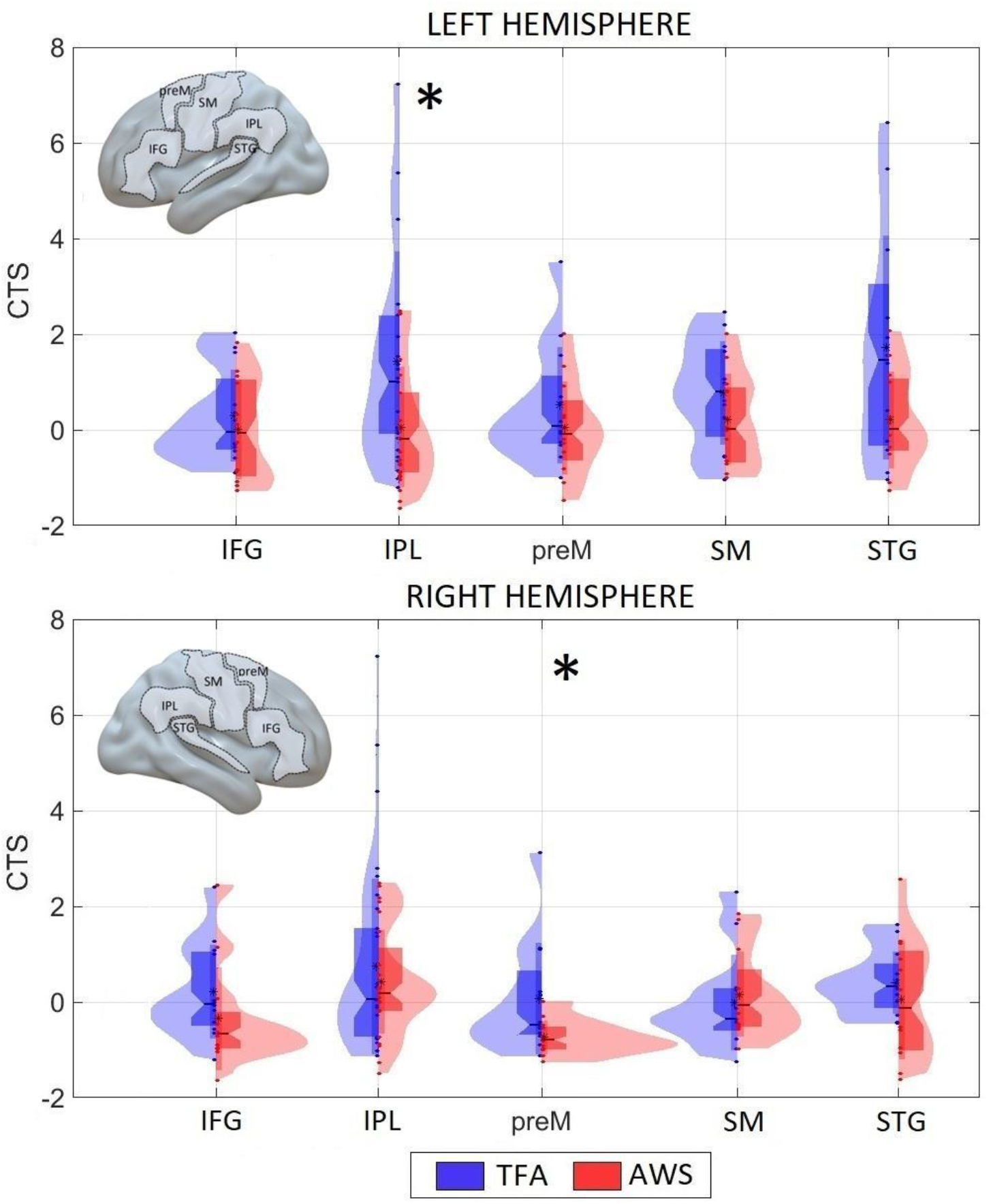
The mean CTS in the 3-5 Hz range (theta band) in each region of interest for each group. Boxplots are overlaid with individual data points and density distributions. Each dot represents data from one of the participants (blue for TFA and red for AWS). Boxes cover the 25th to 75th percentile (inter-quartile range; IQR). The middle of the box represents the median. Whiskers extend from the 25th percentile and 75th percentile to cover all data points lying within 1.5 times the IQR (from the 25th and 75th percentile, respectively). Regions showing a significant group effect are marked with an asterisk.

### Source-level connectivity

We used partial directed coherence (PDC) to assess causal functional connectivity during speech processing in the theta band (3 - 5 Hz) between different ROIs. For the listening-only task (Figure 7), we observed significantly higher connectivity in TFA compared to AWS from the left STG to the right IFG (MTFA = 0.12, SDTFA = 0.05; MAWS = 0.07, SDAWS = 0.04; *p* = 0.01), and from the right IFG to the left IPL (MTFA = 0.03, SDTFA = 0.01; MAWS = 0.02, SDAWS < 0.01; *p* = 0.04). For the listening-for-speaking task (Figure 5), we observed significantly higher connectivity in TFA compared to AWS from the right STG to the left IPL (MTFA = 0.05, SDTFA = 0.06; MAWS = 0.03, SDAWS = 0.02; *p* = 0.05) and from the right STG to the left SM regions (MTFA = 0.11, SDTFA = 0.06; MAWS = 0.06, SDAWS = 0.04; *p* = 0.03).

**Figure 7.**
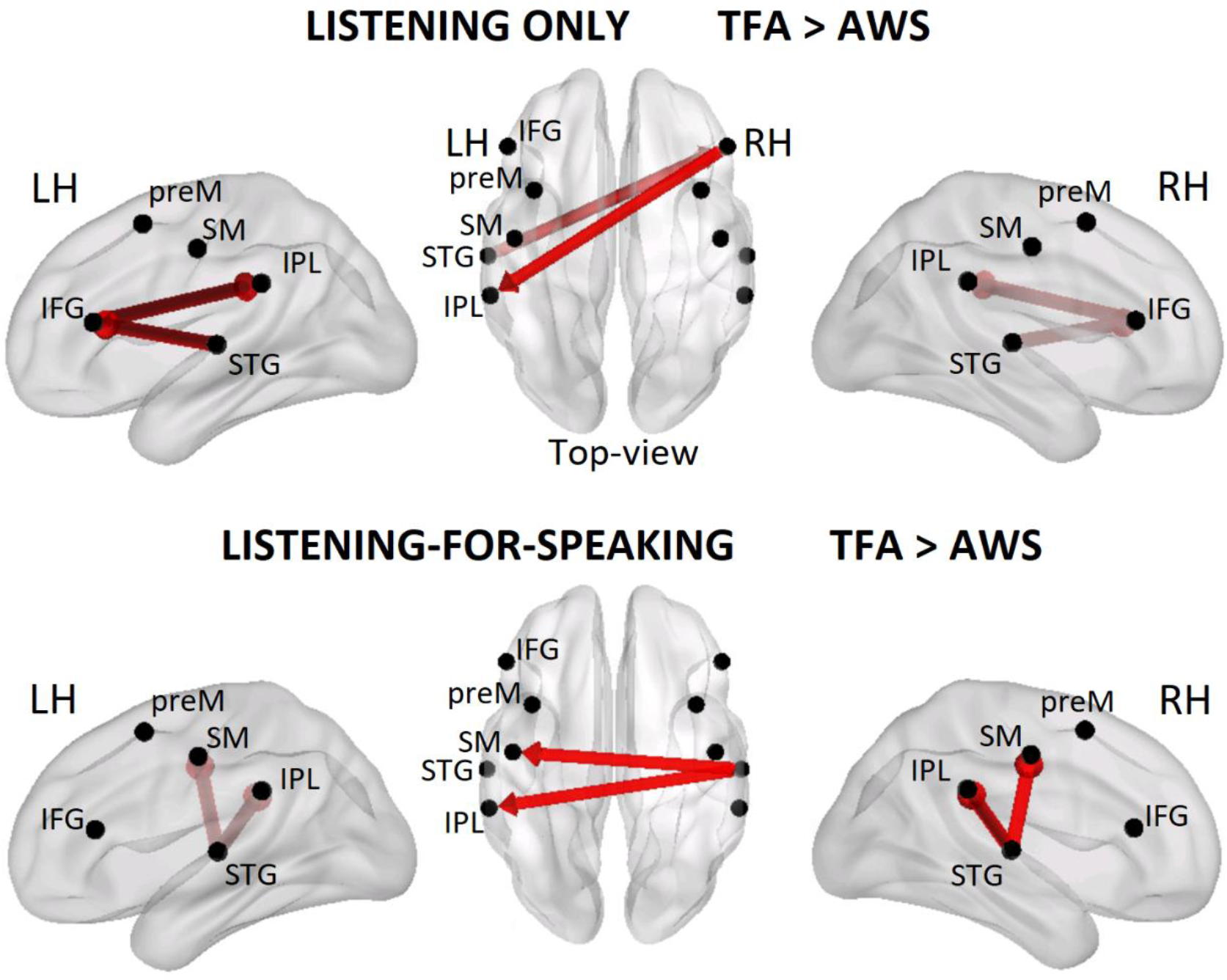
Causal functional connectivity analysis. For each task, we included the connections that exhibited statistically significantly higher PDC values for TFA compared to AWS. We included a seed for each of the regions of interest (IFG: inferior frontal gyrus; preM: premotor/supplementary motor; SM: sensorimotor; IPL: inferior parietal lobule; STG: superior temporal) in both the left (LH) and right (RH) hemisphere.

Interestingly, when considering both groups together, a statistically significant negative correlation was found in the listening-for-speaking task between RTs and the connectivity from the right STG to the left SM cortex (r = -0.56, p = 0.0048): stronger directional connectivity between these regions is associated with faster response times (see Figure 8).

**Figure 8.**
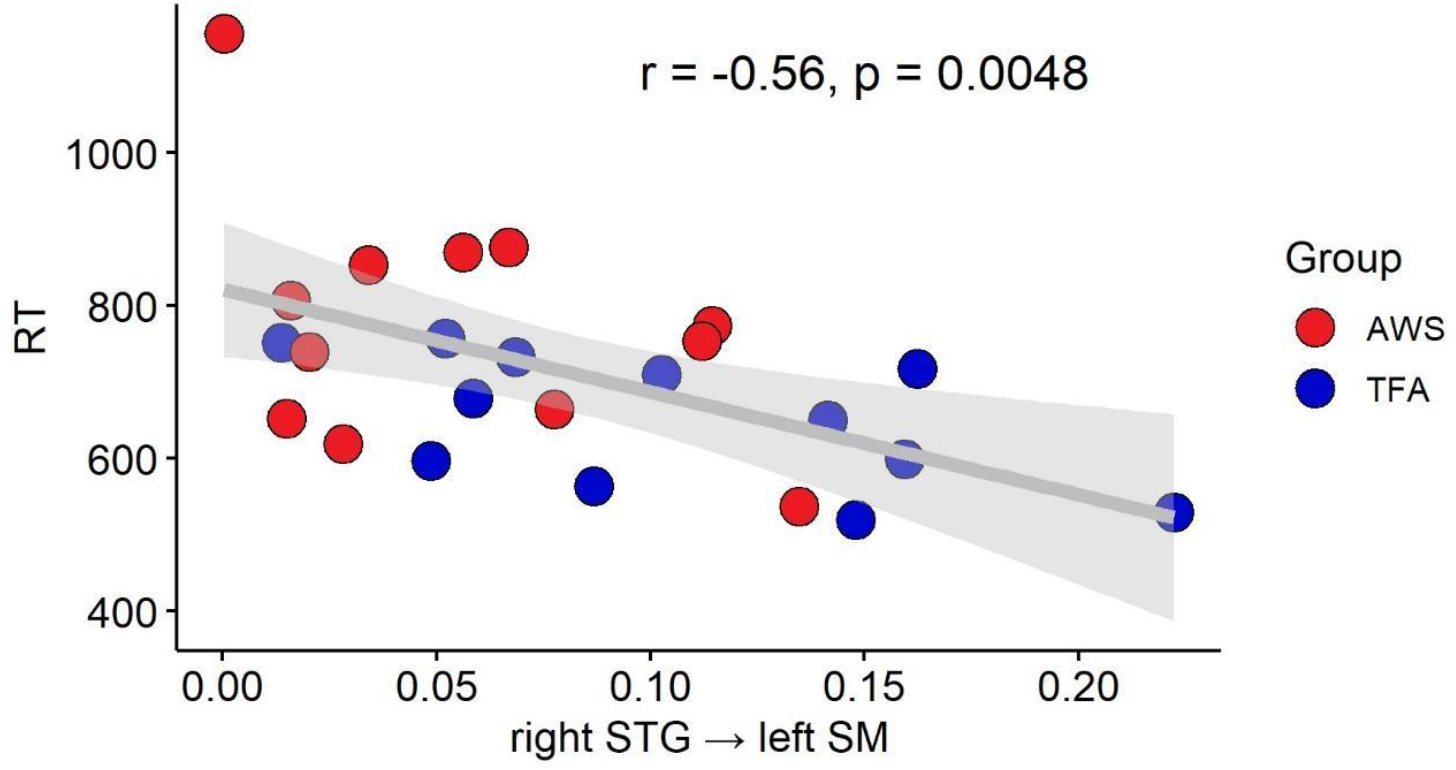
Connectivity-RTs correlation (listening for speaking). Scatterplot showing the correlation between response times (RTs) and connectivity from the right superior temporal gyrus to the left sensorimotor cortex.

## Discussion

In the present work we analyzed cortical tracking of speech (CTS) in a group of participants with dysfunctional speech-motor control due to a neurodevelopmental disorder, namely Developmental Stuttering (DS), comparing adults who stutter (AWS) with typically fluent adults (TFA). To investigate the role of the alertness state of the speech-motor system in CTS, we analyzed two different listening situations: listening-only (no upcoming involvement of speech production) and listening-for-speaking (listen to an unfinished sentence and complete it by naming a picture; upcoming overt engagement of the speech-motor system). We observed reduced coherence in the theta range (3-5 Hz) in AWS relative to TFA in the listening-for-speaking task, both at the sensor (bilaterally around the temporal regions) and the neural source levels. More specifically, at source level, AWS showed lower CTS in the left inferior parietal/temporo-parietal cortex and in the right premotor and supplementary motor regions. Cortical connectivity measures in the theta range were differently modulated as well, with weaker connections in both listening conditions, primarily resulting in lower inter-hemispheric information exchange involving frontal, auditory/temporal and sensorimotor regions. Notably, in the listening-for-speaking task, we also found slower (speech) response times in AWS, and a significant negative correlation between RTs and connectivity from the right STG to the left SM cortex when considering all participants, reinforcing the arguments we lay out next.

### Cortical tracking of speech at the syllabic rate is reduced in Developmental Stuttering when listening for speaking

The listening for speaking condition in this study required speech listening to be interwoven with speech production, similarly to turn-taking in conversational settings (Levinson, 2016). This entails the ability to efficiently time the transition between listening and speaking, and appropriately plan production initiation while still attending to speech. The present findings suggest that CTS in AWS is impaired especially in such situations. As highlighted in the Introduction, CTS is a neural index reflecting the alignment of the phase of (internal) brain frequencies to acoustic features of the speech signal (Assaneo & Poeppel, 2018; Poeppel & Assaneo, 2020; Poeppel & Teng, 2020). Crucially, an intrinsic coupling between auditory and speech-motor regions in a restricted frequency range within the theta band seems to support this process, specifically for the tracking of syllabic rhythm (Assaneo & Poeppel, 2018; Keitel et al., 2018; Morillon & Baillet, 2017; Park et al., 2015). In this study we found that in a population characterized by inefficient timing and implementation of speech-motor processes, i.e., adults who stutter (Alm, 2004, 2021b; Busan, 2020; Chang & Guenther, 2020), CTS is also affected as a result of such intrinsic auditory-motor coupling.

At the source level, in the listening-for-speaking task, we observed CTS reduction in the left inferior parietal cortex and in the right premotor and supplementary motor regions in AWS compared to TFA. All these regions are key cortical substrates for speech-motor coordination. The inferior parietal lobule (IPL), comprising the supramarginal gyrus and the angular gyrus, has been associated with a variety of functions, including verbal working memory, auditory spatial localization, sensorimotor integration, semantic processing and action-motor control (Binder et al., 2009; Binkofski & Buccino, 2018; Bzdok et al., 2016; Shum et al., 2011). Importantly, this region and the partially overlapping (non-anatomically defined) temporo-parietal junction (TPJ; Igelström & Graziano, 2017) are nodes in many dual-route models of speech and auditory processing (Friederici, 2012; Hickok et al., 2011; Hickok & Poeppel, 2004, 2007; Rauschecker, 2012). For instance, in Hickock and Poeppel’s model, the Sylvian Parietal Temporal (Spt) area (located between the inferior parietal lobule and the posterior part of the superior temporal gyrus, thus situated within the TPJ) is proposed to be an interface between auditory codes and motor programs supporting successful sensorimotor integration during speech production, instantiated in the dorsal pathway. The IPL is also key in neurocomputational models of speech production such as the DIVA/GODIVA models (Guenther, 2016), which proposes that somatosensory error maps of the difference between intended and actual somatic states are computed in the IPL during speech production. Importantly, in the adjacent posterior STG/TPJ, auditory error maps are computed by comparing auditory feedback and predicted targets via motor efference copies (Guenther, 2016). Interestingly, even if not properly part of the “classical” cortico-basal-thalamo-cortical network involved in DS (Alm, 2004; Busan, 2020; Chang & Guenther, 2020; Craig-McQuaide et al., 2014), it is not uncommon for this region to be highlighted as part of a defective system in the brain of people who stutter (Busan et al., 2019; Neef et al., 2015; Yang et al., 2016).

On the other hand, the premotor ROI in our study encompasses the premotor cortex and the supplementary motor complex (supplementary motor area – SMA – and pre-SMA). In the speech-motor control literature, these regions have been highlighted in the composition and the timing of execution of speech-motor command sequences (Alario et al., 2006; Ghosh et al., 2008; Guenther, 2016). More specifically, in the DIVA/GODIVA models, the SMA is responsible for the correct initiation of stored speech motor units, while the pre-SMA represents the global sequential structure of the syllables to be produced. On the other hand, these models propose that right hemisphere premotor regions may be a component of a feedback/control speech-motor (Bohland et al., 2010; Guenther, 2016; Tourville & Guenther, 2011; see Chang & Guenther, 2020; Civier et al., 2013 for a perspective on DS). Notably, rhythm processing seems to be particularly reliant upon such cortical structures (together with subcortical regions), both in the speech and non-speech domains (Cannon & Patel, 2021; Fiveash et al., 2021; Kasdan et al., 2022; Ladányi et al, 2020). Additionally, the SMA has also been linked to the mediation of motor-sound representations in auditory prediction and speech imagery (Lima et al., 2016). Crucially, premotor and supplementary motor regions are among the regions that are found to be most dysfunctional in DS (Busan, 2020; Busan et al., 2019; Chang & Guenther, 2020; Civier et al., 2013; A. C. Etchell et al., 2018). When related with the present findings, this body of evidence is compatible with a key role of premotor/supplementary motor regions in tracking rhythmic information in speech perception, particularly at the syllabic level, speculatively by transforming motor information into auditory templates for syllabic tracking.

Given this picture, our results suggest that when upcoming speech is required and neural structures supporting aspects of speech-motor production (i.e., speech-motor sequencing and initiation, rhythmic processing and motor-to-auditory transformation) are inherently inefficient or hinder the proper function of the neural circuit in which they are recruited, as is the case of DS, such structures cannot properly contribute to tracking the speech signal. The fact that we identified regions that are commonly associated with a dorsal stream of speech processing (Friederici, 2012; Hickok & Poeppel, 2007) strengthens the connection between speech-motor abilities and auditory tracking via bidirectional motor-auditory mapping.

The fact that we found differences in CTS within the theta range is particularly interesting from a speech-motor impairment point of view. The theta rhythm has often been associated with syllabic grouping across languages, more specifically to the acoustic energy fluctuations of speech sound clusters organized around an energy peak (usually a vowel) (Strauß & Schwartz, 2017; see also Molinaro & Lizarazu, 2018; Poeppel & Assaneo, 2020). Indeed, a peak was present in the theta range in our audio stimuli (3-4 Hz), reflecting this acoustic property (see Supplementary Figure 2). Importantly, the syllable has been proposed to be an “interface” between the perceptual and the articulatory systems (Poeppel & Assaneo, 2020; Strauß & Schwartz, 2017). Articulators are biomechanically constrained as to the possible configurations they can produce and the speed at which they can be executed; the syllable represents the optimal motor-programming unit that the neural system can send to the motor system for execution (Guenther, 2016; Poeppel & Assaneo, 2020). Crucially, individuals with DS seems to be impaired in the ability to automatically activate syllabic motor units associated with learned sound sequences via the basal ganglia motor loop connected to the pre-SMA and SMA (Alm, 2004, 2021a, 2021b; Busan, 2020; Chang & Guenther, 2020; Civier et al., 2013).

Therefore, it appears that there is a circular relationship that, stemming from biomechanical articulation constraints, via neural motor program units, leads to the acoustic - and hence perceptual - phenomenon of syllabic rhythm tracking (Poeppel & Assaneo, 2020; Strauß & Schwartz, 2017). We believe that this proposed circle of joint causes is strictly related to the results obtained in the present study: the speech-motor production system is involved in tracking acoustic properties that themselves arise from articulatory-motor constraints; when such a system is unstable (as in the case of DS), perceptual tracking is also less efficient, more noticeably when listening and speaking are interwoven and partially overlapping neural resources are required, thus overburdening an already unstable system. Behaviorally, the presence of slower RTs in AWS further supports this view.

### Weaker inter-hemispheric connectivity among bilateral auditory and sensorimotor regions in developmental stuttering when listening to speech

We found weaker connectivity patterns in AWS relative to in both listening conditions. During the listening-only task, we found weaker directional connectivity from the left STG (auditory regions) to the right IFG and from the right IFG toward the left IPL/TPJ. In the listening-for-speaking task, we found weaker directional connectivity from the right STG to the left primary sensorimotor (SM) regions and to the left IPL/TPJ. While we do not interpret the different patterns across listening conditions, all the regions involved are consistent with a dorsal stream of processing (Friederici, 2012; Hickok & Poeppel, 2007), supporting the idea that auditory-motor mapping is important for cortical tracking of speech, at least of syllabic rhythm; this may be related to the nature of the syllable itself, representing the optimal motor unit for the human speech-motor system (Poeppel & Assaneo, 2020; Strauß & Schwartz, 2017). Reduced connectivity in AWS likely reflects lower availability of neural resources for information exchange between regions that are instrumental for auditory and motor processing and integration, compatible with recent proposals suggesting the presence of a general metabolic deficit in the stuttering brain (Alm, 2021a; see also Busan et al., 2019; Han et al., 2019; Maguire et al., 2021; Turk et al., 2021).

Importantly, in the listening-for-speaking task, we found a significant negative correlation between RTs and strength of right STG → left SM cortex connectivity: faster RTs were associated with increased connectivity between these regions. This may indicate that efficiently sending rhythmic auditory information to the primary sensorimotor cortex when speech listening has to be managed with (overt) upcoming speech-motor engagement facilitates speech production, possibly as a result of more efficient CTS and smoother transitioning between listening and speaking with concomitant speech planning. Note that this correlation, when explored separately for AWS and TFA, was not strongly evident in AWS (r = -0.44, p = 0.15) but was present in TFA (r = -0.58, p = 0.045; see Supplementary Table 1). As a further indication, we would also like to highlight that, albeit statistically not significant, an interesting trend was present in AWS when looking at this very same connectivity pattern and SSI-4, where a negative relation is found (r = -0.55, p = 0.06; see Supplementary Table 1 and Supplementary Figure 3): higher SSI-4 scores - hence, more severe stuttering - were associated with weaker right STG → left SM cortex connectivity.

Together, these findings strongly suggest that stuttering may be associated with weaker connectivity between auditory and sensorimotor regions in a preferential frequency range within the theta band (compatible with Assaneo & Poeppel, 2018), which is fundamental for cortical tracking of syllabic units, in turn leading to poorer behavioral performance in terms of response times. This interpretation may be also compatible with proposals according to which an effective connection between these regions may be helpful for better managing (or “by-passing”) disfluencies, perhaps by exploiting rhythmic or tracked cues (A. C. Etchell et al., 2014). Consistent with this suggestive although marginal evidence, it is important to note that right-hemispheric processes, especially in fronto-temporal regions, are often reported as neural markers of stuttering trait and state (e.g., Belyk et al., 2015, 2017; Brown et al., 2005; Budde et al., 2014; Craig-McQuaide et al., 2014; Etchell et al., 2014, 2018; Ingham et al., 2012; Neef et al., 2015; Stasak et al., 2021), suggesting that they may have a role in compensatory (as well as in pathological) speech-motor programming and execution processes in AWS (Busan et al., 2019; Etchell et al., 2014; Neef et al., 2015, 2016, 2018b, 2023).

### Significance of present outcomes for CTS and DS research

The present findings may advance research on both CTS and DS. More specifically, they suggest that 1) CTS requires neural resources that sustain sensorimotor processes for facilitating speech perception and intelligibility, 2) DS may lead to suboptimal CTS, especially when additional resources are needed for supporting concomitant speech preparation for upcoming production, and 3) DS not only impairs speech programming and production but is a more complex neurodevelopmental disorder. Further research should clarify the extent to which DS impacts CTS (and vice versa), how this might affect people’s everyday life, hence widening the scope of possible interventions for stuttering. This is especially important also in light of recent evidence suggesting that auditory-motor coupling (and individual speech production rates) may explain performance in speech comprehension tasks (Lubinus et al., 2023). Speculatively, less efficient CTS may be related to more effortful spoken language comprehension at a subtle level. This is in line with the results reported in Gastaldon et al. (2023): AWS seem less efficient at generating predictions during listening, hypothesized as a result of the inability to fully exploit their speech-motor network. Thus, further studies should investigate whether there is a causal link between CTS and specific processes of speech comprehension such as prediction, and how this causal chain may impact people with different speech and language deficits, especially in interactive contexts (see also Gastaldon et al., 2024 on the importance of studying atypical populations for a better understanding of predictive speech processing). In conclusion, research should move towards turn-based and conversational contexts (e.g., Jackson et al., 2021; Weiss, 1995) to investigate possible subtle differences in how spoken language comprehension is achieved in the stuttering brain.

### Limitations

The study provides interesting new results, suggesting future venues for CTS and DS research; however, some limitations need to be taken into consideration.

First of all, sample sizes are small. The primary reason lies in the difficulty in recruiting AWS participants. This is a common problem when studying neurodevelopmental disorders at low incidence in the population, such as DS (Jones et al., 2002). To address this, in line with increasingly relevant Open Science practices, multi-lab projects can be an efficient way to overcome small *N*’s and to generalize or disconfirm results from individual underpowered studies, and to appropriately quantify effect sizes by means of meta-analyses (Heinrich & Knight, 2020; Lange, 2020; McShane et al., 2019). Note that, by making data available, we provide material for future meta-analyses and/or re-analyses, in the spirit of Open Science.

Another limitation related to DS that should be addressed in future research is that the current study involved male participants only. Persistent DS in adulthood is highly asymmetric according to sex, with a stronger incidence in males (about 1:5 ratio; Yairi & Ambrose, 2013), making recruitment inherently unbalanced. Furthermore, sex hormones may underlie neural changes related to speech-motor control relevant for the persistence or resolution of DS in adulthood (see Neef & Chang, 2024). Thus, it would be interesting to investigate sex-related differences in neural tracking of speech in DS.

Another limitation regards localization of cortical regions. This limitation is common to all studies employing EEG. However, good estimates can still be obtained when using a sufficient number of electrodes covering all the scalp (such as 64 electrodes in the present work), by following standardized electrode placement and by imposing reliable biophysical constraints to forward and inverse solutions (Lantz et al., 2003; Michel & Brunet, 2019; Michel et al., 2004; Westner et al., 2022). Future studies may employ higher density EEG systems or MEG, combined with individual structural scans, in order to provide a more accurate picture.

### Conclusions

The present work suggests that CTS recruits (pre-)motor regions and regions responsible for sensorimotor integration, as well as auditory regions, supporting views proposing an interaction between these networks in speech/language perception (Pickering & Garrod, 2013; Skipper et al., 2017), in addition to their instrumental role in orchestrating successful speech production (Guenther, 2016; Hickok et al., 2011). CTS seems to work less efficiently in DS, especially when additional neural resources are needed for managing listening-for-speaking conditions, as usually happens in more ecological communicative situations (Neef & Chang, 2024). A better understanding of CTS processes in DS under various circumstances may be informative for improving rehabilitation solutions for stuttering.

## Conflicts of interest

None.

## CRediT statement

**Simone Gastaldon:** Conceptualization, Methodology, Formal Analysis, Investigation, Writing – Original Draft, Writing – Review & Editing, Visualization, Project Administration.

**Pierpaolo Busan:** Conceptualization, Writing – Original Draft, Writing – Review & Editing.

**Nicola Molinaro:** Conceptualization, Methodology, Writing – Review & Editing, Supervision.

**Mikel Lizarazu:** Conceptualization, Methodology, Formal Analysis, Data Curation, Writing – Original Draft, Writing – Review & Editing, Visualization, Project Administration.

## Acknowledgments

Research was supported by the University of Padova, by the Basque Government through the BERC 2018–2021 program, and by the Spanish State Research Agency through BCBL’s Severo Ochoa excellence accreditation CEX2020-001010-S. SG was supported by a postdoctoral research grant funded by the Fondazione CARIPARO through the PHD@UNIPD call at the University of Padova (grant CUP_C93C23003190005). NM was supported by the Spanish Ministry of Science, Innovation and University (grants RTI2018-096311-B-I00, PDC2022-133917-I00, PCI2022-135031-2, PID2022-136991NB-I00), the Agencia Estatal de Investigación (AEI), the Fondo Europeo de Desarrollo Regional (FEDER). ML was supported by the Ramón y Cajal programme of Spanish Ministry of Science and Universities (MICIU) (grant RYC2022-035497-I). Funding sources had no role in the collection, analysis, and interpretation of data. We thank Caroline Handley for proofreading the manuscript.

## Data Availability Statement

Data and scripts for the main analyses are available at the OSF repository: https://osf.io/7gpyb/.

## Notes

https://osf.io/7gpyb/

### Competing Interest Statement

The authors have declared no competing interest.

### Summary of Updates

Following peer-review, we revised the manuscript. The Introduction states more clearly what is theoretically-driven and what is exploratory in our study. The Materials and Methods section describes more clearly the coherence computation and substitutes barplots with boxplots in Figure 3A. The Discussion section has been partially rewritten for clarity purposes, and a Limitation section has been included.

