## Supplementary material for "Cortical tracking of speech is reduced in adults who stutter when listening for speaking"

**Content:**

**Supplementary Figure 1**
**Supplementary Figure 2**
**Supplementary Figure 3**
**Supplementary Table 1**

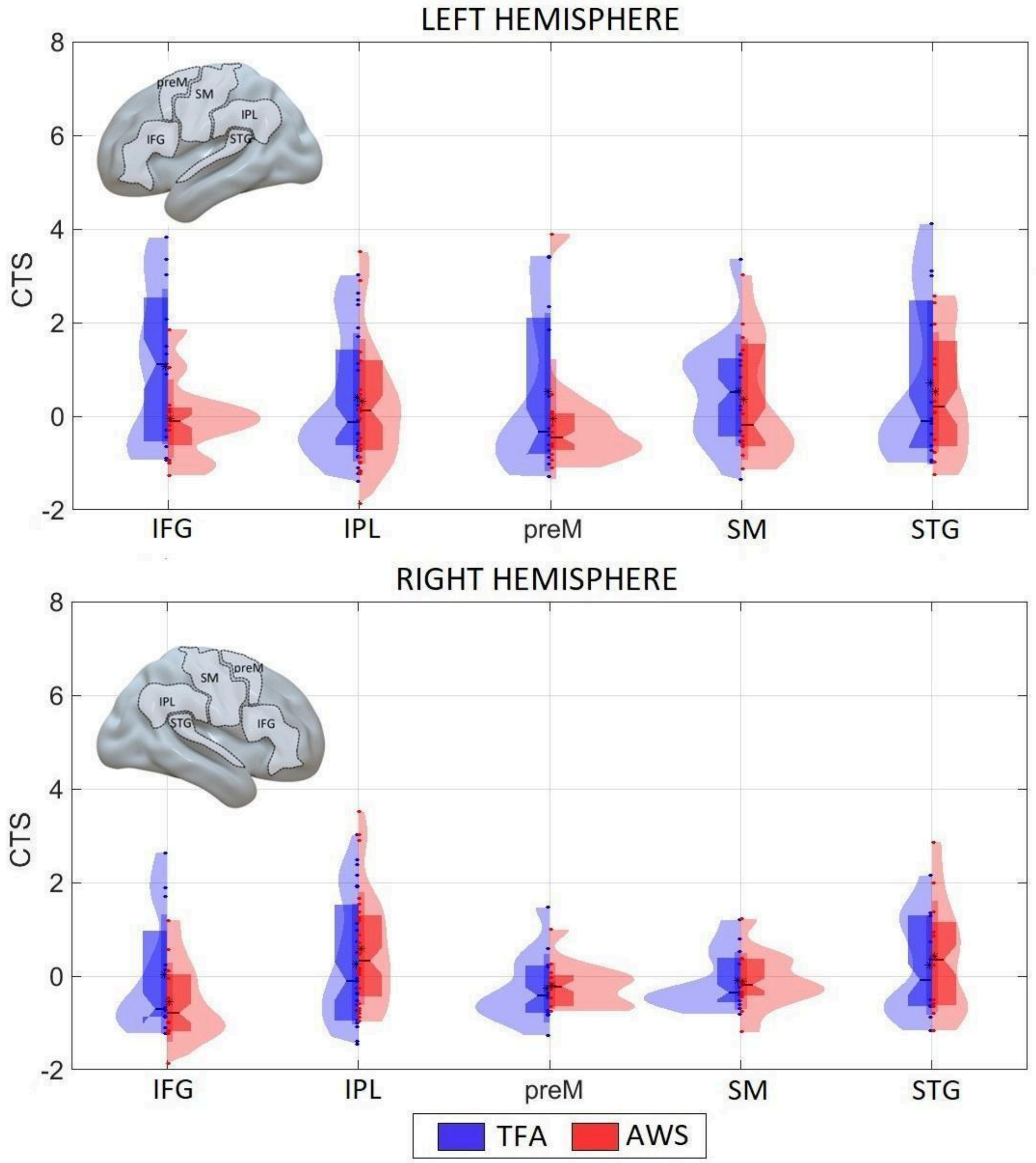


**Supplementary Figure 1:** Source-level CTS in the listening-only task. No statistically significant differences were found.

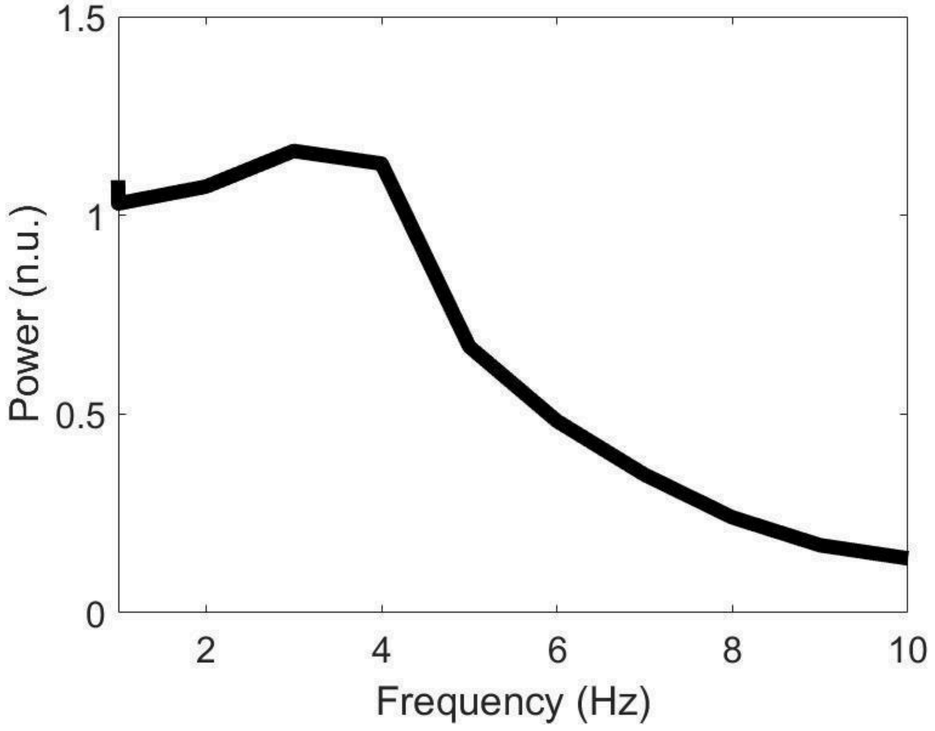


**Supplementary Figure 2:** Spectral characteristics of the speech envelope. We computed the power spectrum of the speech envelope for the frequencies that phase-synchronized with the electrophysiological brain activity (i.e., <10 Hz). The more prominent amplitude modulations were between 3 and 4 Hz, as expected from previous studies (Park et al., 2015; Lizarazu et al., 2019).

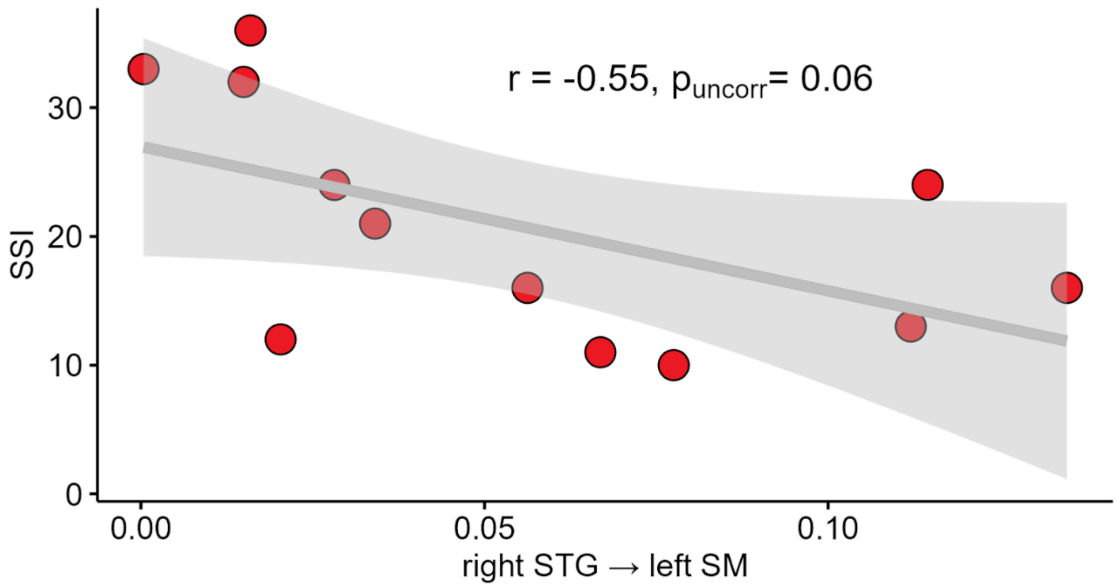


**Supplementary Figure 3.** Scatterplot of the correlation between right STG → left SM and SSI in AWS. Although statistically not significant, a negative trend is observable whereby reduced connectivity from the right STG to the left SM cortex is associated with higher SSI-4 scores (Stuttering Severity Instrument 4; Riley, 2009).

| | ALL PARTICIPANTS | | AWS | | | | TFA | |
|---|---|---|---|---|---|---|---|---|
| | RT | | RT | | SSI | | RT | |
| | r | p | r | p | r | p | r | p |
| SENSOR_COH_LEFT | -0.120 | 0.576 | 0.101 | 0.756 | -0.504 | 0.095 | 0.068 | 0.834 |
| SENSOR_COH_RIGHT | -0.155 | 0.470 | -0.140 | 0.664 | -0.508 | 0.092 | 0.177 | 0.582 |
| SOURCE_COH_L_IPL | -0.038 | 0.859 | -0.051 | 0.876 | -0.272 | 0.392 | 0.458 | 0.134 |
| SOURCE_COH_R_PREM | -0.083 | 0.699 | 0.136 | 0.674 | -0.386 | 0.215 | 0.226 | 0.479 |
| CONN_R_STG_L_IPL | -0.199 | 0.350 | 0.072 | 0.824 | -0.444 | 0.149 | -0.154 | 0.632 |
| CONN_R_STG_L_SM | **-0.556** | **0.005** | -0.439 | 0.153 | -0.551 | 0.064 | **-0.576** | **0.050** |

**Supplementary Table 1:** Correlations between the neural measures found to be different across groups in the listening-for-speaking task (coherence at sensor and source level, connectivity at theta) and behavioral measures (response times and SSI-4). Separate correlations for AWS and TFA are reported only as exploratory.
